# Chiron3D: an interpretable deep learning framework for understanding the DNA code of chromatin looping

**DOI:** 10.64898/2026.03.20.713211

**Authors:** Sebastian Hönig, Aayush Grover, Piero Neri, Didier Surdez, Valentina Boeva

**Author notes:** Equal contribution (Sebastian Hönig, Aayush Grover).

## Abstract

**Motivation:** Three-dimensional folding of the genome into structures such as chromatin loops is essential for gene regulation. Current experimental methods for mapping these structures, like Hi-C and HiChIP, are labor-intensive and require repeated assays to test hypothesized mutation effects. This motivates the need for predictive approaches that reveal the sequence determinants of chromatin loops.

**Results:** In this work, we present a novel and interpretable computational pipeline for predicting CTCF-mediated chromatin loops. We propose Chiron3D, a DNA-only model trained in a cell-type specific manner to predict CTCF HiChIP contact maps. By leveraging pre-trained embeddings from a foundation model, our approach is competitive with baselines that take CTCF ChIP-seq as additional input, while enabling nucleotide-level attribution to the input DNA sequence. Using our framework, we provide likely mechanistic insights into the physical control of loop dynamics. Specifically, we find that the strength of the loop extrusion anchorage site is largely governed by the amount and binding affinity of CTCF sites at the boundaries. Furthermore, we reveal that loop stability is regulated by the amount of intra-loop CTCF binding sites, where fewer sites within the loop lead to a more stable domain. Using targeted, single-nucleotide edit simulations with Chiron3D, we show that both loop strength and stability can be precisely controlled. Together, these results provide novel mechanistic insights into the physical control of genome organization and highlight the potential of decoding the DNA sequence logic in silico.

**Availability:** The Chiron3D pipeline is made available at https://github.com/BoevaLab/Chiron3D.

**Supp. information:** Supp. data are available at Journal Name online.

## Introduction

Three-dimensional genome folding brings distal regulatory elements into close proximity to regulate gene expression. Topologically Associating Domains (TADs) are primarily formed by loop extrusion (3), a process in which cohesin translocates along chromatin and progressively enlarges loops until it is halted by CTCF boundary elements (14). Genome-wide conformation assays such as Hi-C (11), Micro-C (8), and HiChIP (13) map these interactions across the genome with high resolution or protein specificity. Extrusion appears as stripes intersecting loop anchors; these are enriched for inward-oriented CTCF motifs, and disrupting them abolishes striping patterns (26). Loop domains also display pronounced corner interactions, where boundary-to-boundary contacts are much stronger than along the stripe (5). Reduced striping with strong corner enrichment indicates extrusion constrained near boundaries, enhancing structural stability.

Building on these insights, several deep learning models predict 3D genome folding from DNA sequence. DeepC and Akita predict locus-specific contact maps from 1 Mb of sequence and estimate variant effects (20; 2), while Orca extends this to whole chromosomes, capturing long-range compartmentalization (29). A recent work has explored the sequence grammar underlying genome folding, showing that deep learning models can reveal how CTCF motif combinations determine loop architecture (22). For cell type specificity, C.Origami integrates DNA with ATAC-seq and CTCF ChIP-seq to predict Hi-C maps and enables in silico perturbation screening (24), whereas EPCOT and UniversalEPI use DNA and ATAC-seq to generalize enhancer–promoter interactions across unseen cell types (28; 4). Nonetheless, there is a need for a DNA-only model that performs competitively in predicting HiChIP contact maps while preserving nucleotide-level interpretability to study sequence-encoded determinants of loop stability.

CTCF HiChIP offers an enriched view of extrusion-stabilized loops, making it a strong target for learning how DNA sequence encodes loop extrusion. Since alterations at single nucleotides can modify the affinity of CTCF binding and disrupt boundary elements, a sequence-only framework offers unique advantages for assessing how such changes reshape local chromatin topology. Motivated by this, we present Chiron3D, a DNA-only attention model initialized with Borzoi embeddings (12) designed to predict CTCF HiChIP contact maps within approximately 524 kb windows. Furthermore, we introduce a training and analysis pipeline enabling interpretation of individual loop loci using two complementary approaches: (i) TF-MoDISco (21) for motif discovery from attribution scores and (ii) an edit framework based on Ledidi (19) to propose sparse nucleotide edits that alter loop type. Chiron3D recapitulates the experimental findings of Vian *et al*. (26), demonstrating that the strength and number of the CTCF binding motifs govern loop anchorage, and show that localized sequence edits can induce stripe formation or invert asymmetry programmatically. Furthermore, by examining the determinants of loop stability, we find that intra-loop occurrences of CTCF binding sites tend to destabilize loop boundaries and this effect can be modulated in silico through few targeted edits along the genome. Overall, Chiron3D approximates cell-type-specific HiChIP structures and enables mechanistic manipulation of sequence features governing stripe formation and loop stability, providing novel insights into the processes underlying chromatin loop formation.

## Methods

Our pipeline for sequence-driven analysis of chromatin looping comprises three modules (Fig. 1): (A) Training of the Chiron3D model, where a DNA-only attention network initialized with Borzoi embeddings (12) is trained to predict cell-specific HiChIP contact maps from 524,288 base pair (bp) one-hot encoded DNA sequences; (B) Motif discovery, which produces nucleotide-level attribution scores for the loop boundary regions and converts these into sequence motifs using TF-MoDISco (21); and (C) Genomic editing, which makes use of the Ledidi framework (19) to propose sparse nucleotide edits that optimize for the defined objectives such as stripe enhancement, stripe reduction, or loop stability.

**Figure 1.**
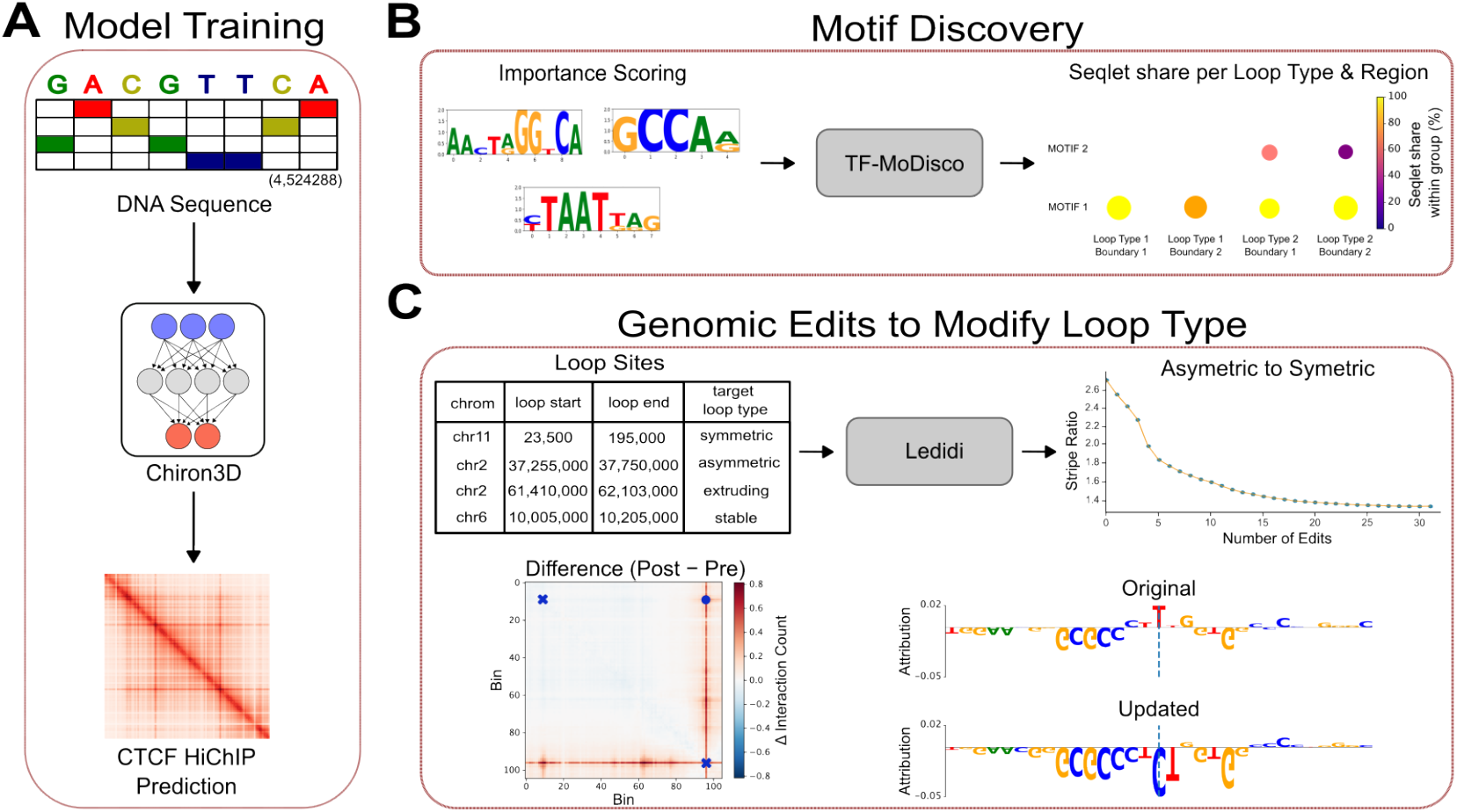
Overview of the proposed pipeline. **A**, Chiron3D is a deep learning model that predicts CTCF HiChIP contact maps from approximately 524 kb DNA sequence. **B**, Using our pipeline including Chiron3D and TF-MoDISco, we can identify transcription motifs important for long-distance chromatin contacts mediated by CTCF binding at loop boundaries. **C**, Loop types can be modified using few single nucleotide edits using our pipeline containing Chiron3D and Ledidi.

### Data

We analyzed CTCF HiChIP together with the matched CTCF ChIP-seq from the A673 wild-type Ewing sarcoma cell line reported by Surdez *et al*. (GEO: GSE133227) (23). The genome was tiled into 524,288 bp windows with a 50 kb stride, chosen to encompass the CTCF-mediated loops (median loop length ≈185 kb (16; 18; 6)).

Each window was mapped to a 105 × 105 target matrix at 5 kb resolution. Windows overlapping chromosome ends or intersecting the hg19 blacklist regions were excluded.

We used two datasets of loop sites. We used Tweed-derived loop annotations from Surdez *et al*. (23), which classifies loops as symmetric (XY) or asymmetric (X or Y). For a loop with boundaries *x*_*L*_ and *x*_*R*_ (*x*_*L*_ *< x*_*R*_) in contact matrix *M*, we excluded the inner *u* = 15 bins (75 kb) near the boundaries and defined the stripe index sets as ℬ _*L*_ = {(*i, x*_*L*_) : *i* = *x*_*L*_ + *u*, …, *x*_*R*_} and ℬ _*R*_ = {(*x*_*R*_, *j*) : *j* = *x*_*L*_, …, *x*_*R*_ − *u*}. The corner interaction is given by *M* (*x*_*L*_, *x*_*R*_) whereas the mean stripe intensities (*S*_*L*_ for stripe along left boundary, *S*_*R*_ for stripe along right boundary) are defined as

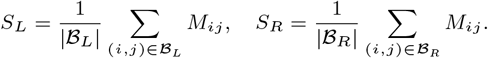

Following Surdez *et al*., loops were labeled asymmetric using stripe ratio 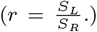 or *r* ≤ *µ* − 2 SD, where *µ* and SD are the mean and standard deviation of *r* over the dataset set and symmetric otherwise. This yielded 8,017 total loops, of which 696 were asymmetric under this criterion and the remaining were symmetric (Supp. Fig. S1).

Stable loops were identified using FitHiChIP, with additional filtering applied to the resulting significant loops (Supp. Methods) (1).

### Chiron3D Architecture

We initialized the sequence backbone using a pretrained Borzoi model (12; 7), which serves as a sequence feature extractor encoding regulatory and epigenomic signals relevant to chromatin organization. Given a one-hot encoded DNA-sequence spanning approximately 524 kb, the backbone produces contextualized sequence embeddings that capture long-range regulatory information (Fig. 2A). These embeddings are processed by a multi-head self-attention module that models interactions across the genomic window. A lightweight U-Net-style decoder (17) then aggregates these representations. Pairwise interaction features are constructed from position-wise embeddings and refined using dilated residual blocks to capture multi-scale chromatin interaction patterns.

**Figure 2.**
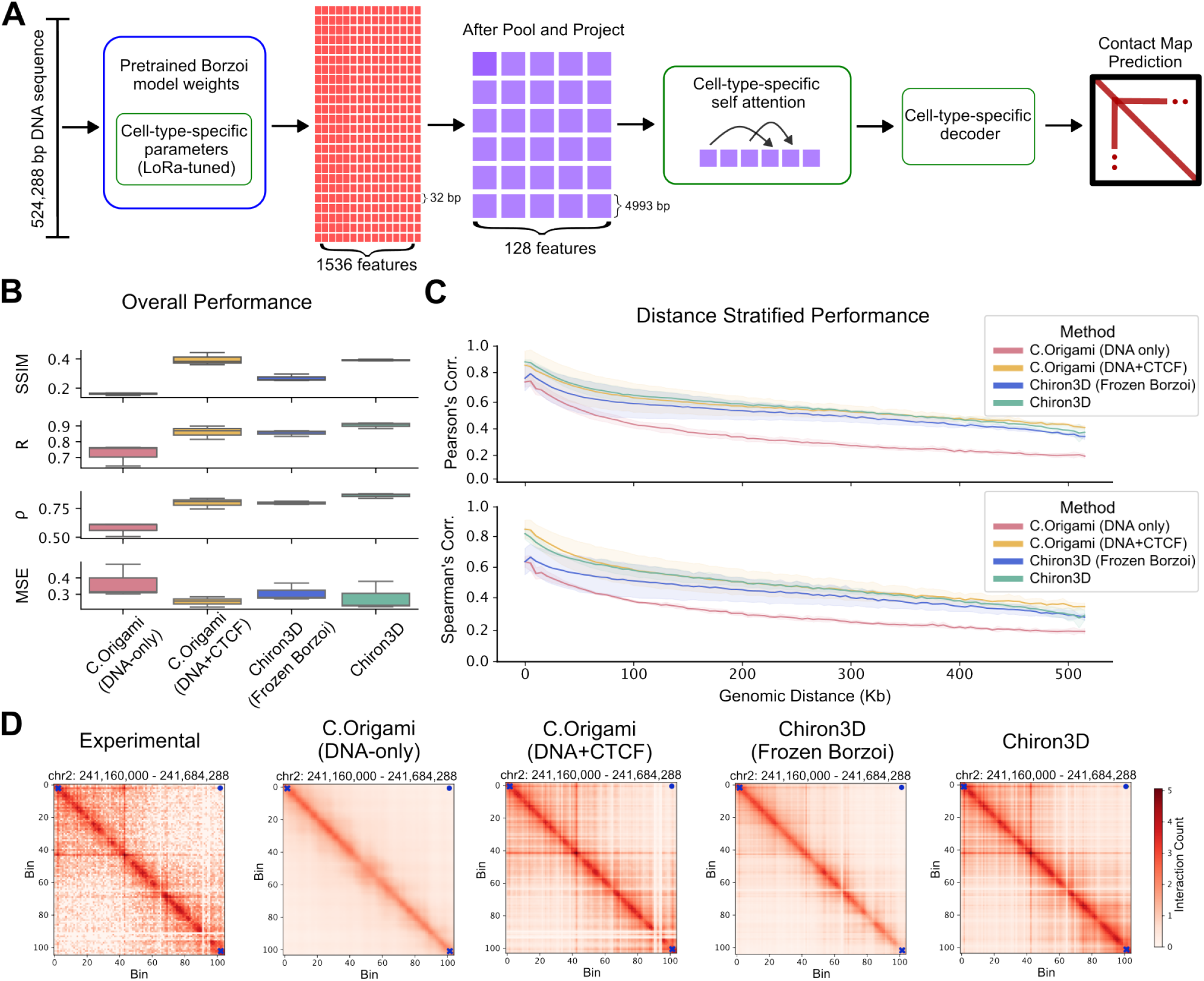
Chiron3D performs comparably to C.Origami, without requiring a CTCF ChIP-seq input. **A**, Detailed architecture of Chiron3D. **B**, Structural Similarity Index Measure (SSIM), Pearson’s correlation (*R*) of the insulation values, Spearman’s correlation (*ρ*) of the insulation values, and mean squared error (MSE), calculated on the test set. Each point represents metric between experimental and predicted contact map for a given test chromosome. **C**, Distance-stratified Pearson’s and Spearman’s correlation of the insulation values binned at 5 kb. Shaded regions show 95% bootstrap confidence intervals. **D**, Qualitative comparison between model predictions and experimental HiChIP at a representative test locus on chromosome 2. The loop boundaries are marked with blue crosses whereas the corner interaction is depicted with a blue dot.

To adapt the pretrained backbone to the HiChIP prediction task, we considered two regimes: (i) a frozen backbone with a trainable prediction head and (ii) parameter-efficient fine-tuning using low-rank adaptation (LoRA) (9). In the latter case, only a small subset of parameters is updated while the majority of the pretrained backbone remains fixed.

Finally, to map the embeddings to the 105 × 105 contact target (525 kb at 5 kb resolution),we utilized a sequence-to-matrix architecture comprising linear projections, transformer blocks (25), and a dilated residual decoder (Fig. 2A, Supp. Fig. S2). Additional implementation and model training details are provided in the Supp. Methods.

### Optimization Objectives

We define quantitative objectives for stripe asymmetry and loop stability, which guide downstream attribution and sequence-edit analyses aimed at altering loop types:

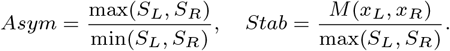

where *S*_*L*_ and *S*_*R*_ denote mean left and right stripe intensities respectively, and *M* (*x*_*L*_, *x*_*R*_) represents the corner interaction between boundaries. *Stab* quantifies the relative strength of the corner interaction compared to the dominant stripe, with higher values indicating stronger anchoring. *Asym* instead quantifies imbalance between stripes, with larger values reflecting greater asymmetry (Supp. Fig. S1).

## Results

### Chiron3D accurately predicts HiChIP contact maps from DNA sequence alone

We trained and evaluated Chiron3D on the A673 cell line from Surdez *et al*. (23), which provides high-quality matched CTCF HiChIP and ChIP-seq datasets. The model predicted log(1+*x*)-transformed HiChIP contact maps at 5 kb resolution from a 524,288 bp DNA input, yielding a 105 × 105 output matrix (Fig. 2A).

We benchmarked Chiron3D against two variants of C.Origami (24), the current state-of-the-art model with comparable resolution. To isolate DNA-sequence–driven predictive performance, we primarily compared Chiron3D with the DNA-only variant of C.Origami. We additionally included a C.Origami model augmented with CTCF ChIP-seq input to provide an upper bound on performance with minimal epigenetic input. For Chiron3D, we evaluated two variants:

(i) a frozen-Borzoi model and (ii) a LoRA-tuned model (9), which fine-tunes a small subset of parameters using low-rank adaptation.

The performance of Chiron3D was assessed using structural similarity index measure (SSIM) (27), mean squared error (MSE), and Pearson’s and Spearman’s correlations of the insulation track on the test set, together with distance-stratified correlations to evaluate performance as a function of genomic separation. C.Origami (DNA-only) performed suboptimally (SSIM 0.16; MSE 0.33; Pearson 0.75; Spearman 0.60; Fig. 2B). Incorporating CTCF ChIP-seq markedly improved its accuracy (SSIM 0.39; MSE 0.25; Pearson 0.88; Spearman 0.82). In contrast, Chiron3D with LoRA achieved the strongest overall performance (SSIM 0.39; MSE 0.25; Pearson 0.91; Spearman 0.87), while the frozen-Borzoi variant showed moderate performance (SSIM 0.27; MSE 0.29; Pearson 0.86; Spearman 0.80). Despite being a DNA-only model, Chiron3D matched the performance of the CTCF-augmented C.Origami model across all genomic distances (Fig. 2C).

To better understand these differences, we examined individual loci from the test set (Fig. 2D, Supp. Fig. S3). All models except C.Origami DNA-only recovered the characteristic striping loop pattern, though Chiron3D with a frozen Borzoi backbone exhibited slightly weaker signal intensity. Overall, Chiron3D performed competitively with the C.Origami model even when C.Origami used CTCF ChIP-seq as an additional input, while outperforming C.Origami DNA-only model. Importantly, Chiron3D requires only DNA sequence as input, preserving nucleotide-level interpretability and providing a foundation for downstream motif analysis.

### Loop symmetry and anchoring are driven by CTCF motif strength and count at boundary regions

Chromatin loops arise through the process of loop extrusion, where cohesin complexes progressively reel in DNA until they encounter boundary elements such as CTCF. Loops can appear symmetric when both boundaries contribute equally to anchoring or asymmetric when one boundary serves as the dominant anchor (Fig. 3A). On contact maps, this distinction manifests as a stripe pattern extending from one or both anchors. We applied the trained Chiron3D model to identify DNA sequence-encoded determinants of loop asymmetry.

**Figure 3.**
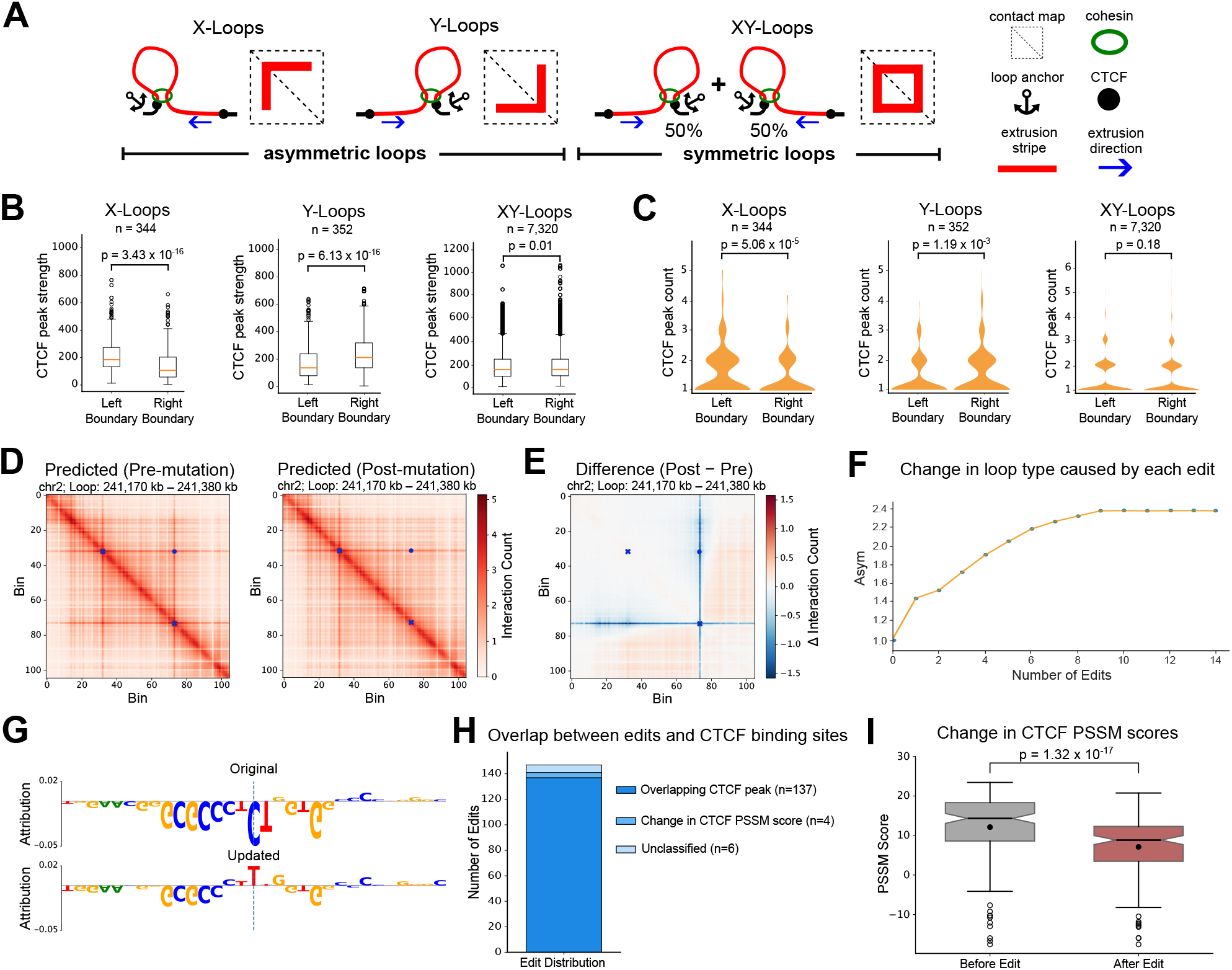
CTCF anchoring strength and count define loop symmetry and stripe dynamics. **A**, Schematic illustration of symmetric XY-loops and asymmetric X- and Y-loops and contact map patterns. **B**, Measurements of CTCF ChIP-seq signal at loop boundaries of X- and Y-asymmetric loops and symmetric loops. *n* is the number of loops of each type. **C**, Number of CTCF ChIP-seq peaks at loop boundaries of X- and y-asymmetric loops and symmetric loops. **D**, Example of a region from the test set where a single edit is proposed to make a symmetric loop more asymmetric. **E**, The difference in predicted contact maps caused by the proposed edit. **F**, Line plot showing change in the optimization function (*Asym*) as a function of number of proposed edits. **G**, Attribution scores before and after the edit was introduced in the sequence. **H**, Proportion of edits, proposed for symmetric loops in the test dataset, that overlap with CTCF regions. **I**, Change in CTCF PSSM scores caused by the edits across all symmetric loops in the test data. All p-values are calculated using Wilcoxon signed-rank test.

To quantify asymmetry, we defined stripes as horizontal or vertical bands extending from loop boundaries. The degree of asymmetry was captured by the *Asym* score where larger values indicate stronger anchoring to a single boundary (Methods). We categorized loops by anchorage: left-anchored as X-loops, right-anchored as Y-loops, and both-sided as symmetric (XY) loops (Supp. Fig. S1).

To examine the molecular basis of this asymmetry, we analyzed CTCF ChIP-seq binding strength at both loop boundaries, taking the maximum signal within each region. This analysis was performed separately for each class of asymmetric loops and for all symmetric loops. Consistent with the previously observed anchoring behavior (26), the experimental CTCF ChIP-seq signal was stronger on the right boundary for Y-loops and on the left boundary for X-loops, corresponding to the loop’s anchor site (Fig. 3B). In contrast, symmetric loops showed no such bias (Fig. 3B). Moreover, similar trend was observed in the number of CTCF ChIP-seq peaks across symmetric and asymmetric loops (Fig. 3C). These results were further confirmed by training Chiron3D on cohesin (SMC1A) HiChIP data (Supp. Fig. S4).

Having characterized asymmetry using experimental data, we next evaluated whether our model, Chiron3D, captures similar sequence-level determinants. We computed importance scores for all loop boundaries using Chiron3D and employed TF-MoDISco across all Tweed-loops (21), stratified by loop classes (Supp. Methods). CTCF emerged as the dominant motif for the *Asym* objective, with no other motif approaching its importance (Supp. Fig. S5). Subsequent analyses therefore focus on CTCF-mediated anchoring.

Next, we tested whether single-nucleotide edits can convert symmetric loops to asymmetric loops, given that single-nucleotide polymorphisms (SNPs) or single-nucleotide variants (SNVs) in cancer can potentially impact 3D genome architecture (10). This was done using the Ledidi model (19) that proposes minimal sequence changes leading to altered stripe balance in predicted HiChIP contact maps (Supp. Methods). We first selected the 86 loop sites from the three test chromosomes with near-symmetric stripe ratio (close to 1) and optimized edits to increase asymmetry. For example, at a 210 kb loop on chr2: 241.17 - 241.38Mb, Ledidi proposed a single substitution (C→T) at position chr2: 241,378,546 within the right boundary’s CTCF peak, reducing the Y stripe signal (Fig. 3D-E). This single edit maximized *Asym* score for this locus (Fig. 3F). The substitution occurred at position 10 of the CTCF motif (Supp. Fig. S6), reducing the motif’s PSSM score from 14.32 to 5.63 and increasing *Asym* from 1.07 to 1.46 (Fig. 3G).

Across all 86 loops, Ledidi proposed 147 edits, 137 of which overlapped with a CTCF ChIP-seq peak (Fig. 3H, Supp. Methods). Of the remaining 10 edits, 4 sites were classified as CTCF by their PSSM score changes and the remaining 6 were unclassified (Fig. 3H). The mean CTCF PSSM score for these edits decreased by 6.29 (Fig. 3I), consistent with the motif disruption leading to the loss of anchoring and increased asymmetry.

Similarly, we reversed the objective on 87 asymmetric loops to equalize stripe strengths. Across these loops, Ledidi proposed 204 edits: 107 overlapping CTCF peaks and 86 additional edits that created or reinforced motifs, increasing mean PSSM by 2.26 (Supp. Fig. S7).

Finally, we examined the positions of all proposed edits within the canonical CTCF motif. Edits strengthening or creating novel CTCF binding sites were evenly distributed across the motif, while disruptive edits primarily targeted nucleotides with largest bit scores (Supp. Fig. S6). Out of the 351 edits proposed by our framework to alter loop anchoring, 26 corresponded to known SNPs in dbSNP (15) (Supp. Table S1), suggesting that our framework can identify naturally occurring variants that may influence chromatin architecture. Overall, these results highlight that Chiron3D, when coupled with Ledidi, learns sequence-level determinants of boundary strength and exploits key nucleotides within the CTCF motif to alter loop anchoring and stripe symmetry.

### Loop stability is subject to the amount of intra-loop CTCF binding sites

In contrast to the prominent striping patterns, stable loops are defined by strong boundary-to-boundary interactions and minimal intra-loop HiChIP signal (Fig 4A). To quantify this, we defined loop stability as the ratio of corner interaction strength to the maximum mean stripe intensity, with higher values indicating greater stability. Stable loops were identified using FitHiChIP, excluding overlaps with Tweed loops (Supp. Methods). Compared to extruding loops, stable loops showed over twofold fewer CTCF ChIP-seq peaks per unit length (Fig. 4B), suggesting a novel relationship between loop stability and reduced intra-loop CTCF density. Since the entire input DNA sequence was of interest and TF-MoDISco is limited in scope, we focused on sequence edits with Ledidi rather than motif discovery to identify relevant transcription factor sites.

**Figure 4.**
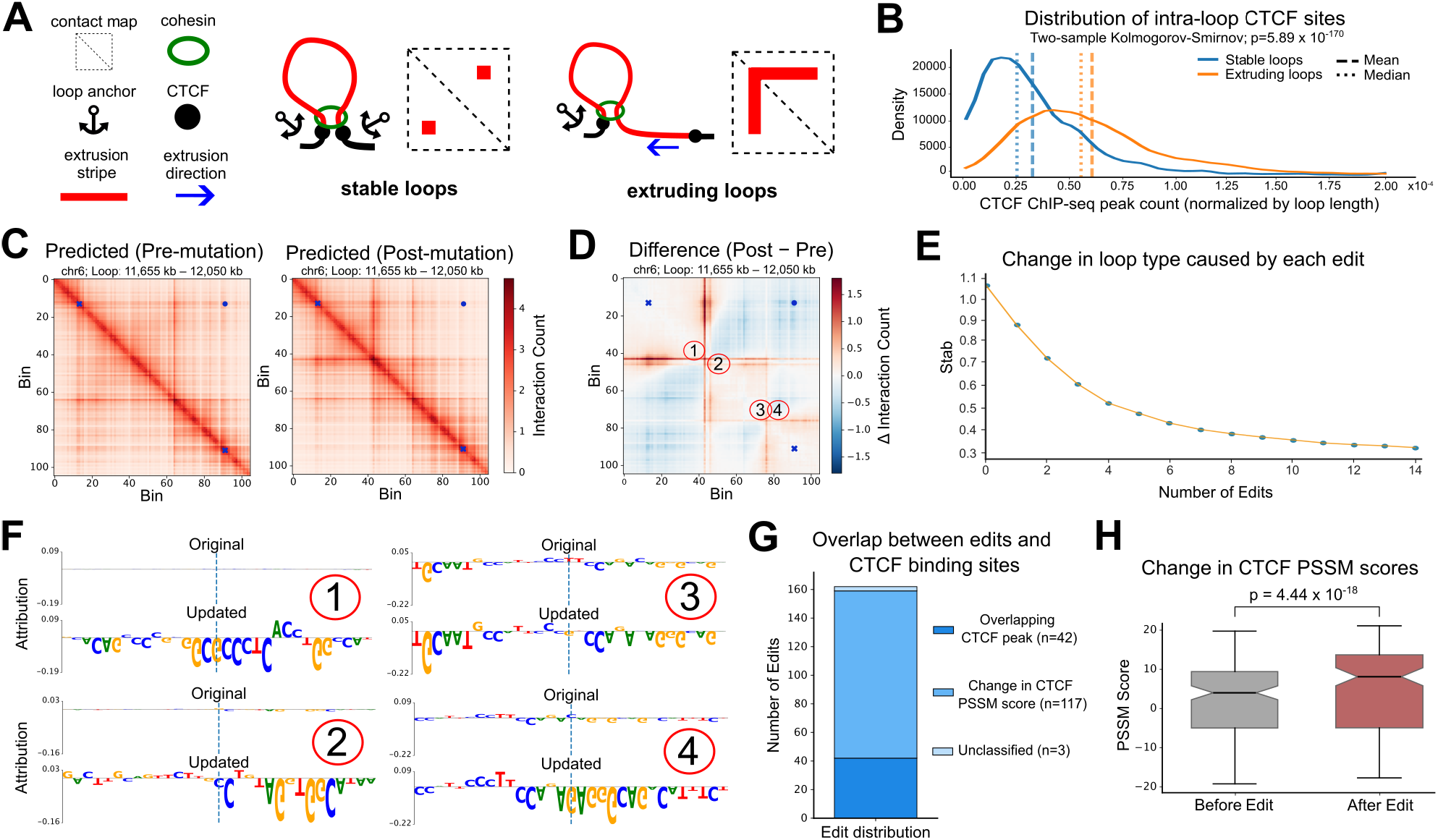
The amount of intra-loop CTCF sites defines loop stability. **A**, Schematic illustration of stable and extruding loops and contact map patterns. **B**, Density plots displaying the distribution of length normalized CTCF peak frequencies for stable and extruding loops. The *p*-value is calculated using two-sample Kolmogorov-Smirnov test. **C**, Example of a region from the test set where four edits are proposed to convert the loop from stable to extruding. **D**, The difference in predicted contact maps caused by the proposed edits. **E**, Line plot showing change in the optimization function (*Stab*) as a function of number of proposed edits. **F**, Attribution scores before and after each edit was introduced in the sequence. **G**, Proportion of edits, proposed for stable loops in the test dataset, that overlap with CTCF regions. **H**, Change in CTCF PSSM scores caused by the edits across all stable loops in the test data. Significance is calculated using Wilcoxon signed-rank test.

After establishing the experimental differences between stable and extruding loops, we applied the Ledidi framework with Chiron3D to test whether sequence edits can tune loop stability. Optimizing for increased extrusion, Ledidi proposed edits that strengthened or created new CTCF binding sites. For example, in a loop on chr6: 11.65 - 12.05 Mb, four edits were proposed that increased intra-loop interactions and decreased the interactions between the boundary regions (Fig. 4C,D). The edits form new internal loops rather than enhancing the primary stripes. These internal loops divert interaction frequency away from the main anchors, heavily depleting the corner signal. Quantitatively, this reduced the stability score (*Stab*) from 1.07 to 0.52 (Fig. 4E). These four edits altered the attribution scores in their respective neighborhoods (Fig. 4F). Across 87 highly stable loops, 162 edits were proposed, with 42 in existing CTCF peaks and 117 introducing new or stronger motifs (Fig. 4G), increasing mean PSSM scores from 2.56 to 5.61 (Fig. 4H).

Conversely, when optimizing for increased stability, Ledidi proposed edits that raised *Stab* in the predicted contact maps. In a representative loop (chr2: 219.89 - 220.27 Mb), five edits within CTCF peaks increased stability from 0.65 to 0.82 (Supp. Fig. S8). Across 75 extruding loops, 247 edits were predicted, of which 244 disrupted CTCF motifs, lowering their PSSM scores by an average of 8.52 (Supp. Fig. S8). For both objectives, edits were restricted to the inter-boundary interval with a boundary-exclusion mask: we disallowed edits at the boundaries and within five bins (25 kb) inward of each boundary. Without this mask, edits cluster at boundaries and can artificially alter *Stab* (Supp. Fig. S9).

Finally, examining all proposed edits within the canonical CTCF motif, we again observed that edits that strengthened or created novel CTCF binding sites were evenly distributed across the motif, whereas disruptive edits primarily targeted nucleotides with largest bit scores (Supp. Fig. S6). Among 409 predicted edits that alter loop stability, 59 coincided with known dbSNP variants (15) (Supp. Table S2). These results indicate that our framework can propose biologically plausible, naturally occurring sequence edits relevant to chromatin organization. Together, these findings suggest that loop stability can be bidirectionally tuned by single-nucleotide edits that alter CTCF occupancy within loop domains.

## Discussion

In this work, we investigated DNA-encoded mechanisms underlying chromatin looping. We analyzed CTCF HiChIP contact maps that capture extrusion-stabilized CTCF-anchored loops and provide a direct readout of loop extrusion dynamics. These data distinguish stable from unstable loops and symmetric from asymmetric configurations.

Our bioinformatic analysis confirmed and quantitatively extended previous observations (26) that stripe anchoring, *i.e*., dominance of a loop boundary in generating extrusion-associated stripes, is associated with higher CTCF motif strength and count at loop boundaries (Fig. 3). Using Chiron3D, a DNA-only attention model for predicting HiChIP contact maps, we further investigated factors governing chromatin loop dynamics at nucleotide resolution (Fig. 1). Chiron3D was benchmarked against the state-of-the-art transformer-based architecture C.Origami (24) and demonstrated comparable accuracy in predicting cell-type-specific contact maps, despite C.Origami incorporating CTCF ChIP-seq as an additional input (Fig. 2). Unlike previous models, such as C.Origami and UniversalEPI (4), which use additional inputs such as ATAC-seq and CTCF ChIP-seq and prioritize predictive accuracy over mechanistic interpretability, Chiron3D enables high-resolution prediction with direct manipulation of sequence determinants governing loop topology. Our model is also substantially lightweight than other DNA-only models, such as Orca and DeepC (29; 20), trained to predict Hi-C maps. By integrating sparse edit simulations with nucleotide-level attribution, our framework enables testing causal hypotheses about chromatin looping that were previously inaccessible without labor-intensive genome editing experiments.

Combining Chiron3D with motif-exploration tools TF-MoDISCo and Ledidi, we linked CTCF motif strength at loop boundaries with striping patterns that characterize loop dynamics. In silico mutagenesis with controller-guided sparse edits demonstrated that targeted sequence perturbations weakening or strengthening CTCF motifs modulate striping patterns and shift loop symmetry in the intended direction.

Furthermore, using our pipeline, we revealed that more stable loops exhibit a depletion of intra-loop CTCF motifs compared to extruding loops (Fig. 4). In silico edits demonstrated that weakening intra-loop CTCF sites increases stability, while strengthening or introducing new sites decreases it, with effects scaling with the number of edits. These results suggest that regulatory variants, including non-coding disease-associated SNPs, may alter 3D genome organization by subtly modulating CTCF motif strength, linking sequence variation to higher-order chromatin structure.

Our findings with the Chiron3D-based pipeline extend prior observations on CTCF motif orientation, strength, and multi-site enrichment at loop anchors by (i) identifying minimal, single-nucleotide edits that drive topological transitions, and (ii) quantifying the loop stability continuum. The DNA-only framework further offers nucleotide-level interpretability not achievable in multimodal models. While all edits were evaluated in silico, we acknowledge that chromatin loop formation is influenced not only by DNA sequence but also by dynamic factors such as transcriptional activity, and cell-cycle state, which are not explicitly modeled here. Additionally, our analysis focused on a single cell line, and extending this framework across diverse cellular contexts will be essential to assess generalizability. Future extensions to additional assays (*e.g*., Hi-C) and cell types will further advance our understanding of the sequence determinants of 3D genome organization.

Overall, Chiron3D establishes a sequence-based, interpretable framework for dissecting DNA-encoded mechanisms of chromatin looping. By enabling targeted edits that modulate loop symmetry and stability, it provides a powerful in silico tool to probe the causal role of nucleotides in 3D genome organization.

## Notes

### Competing Interest Statement

The authors have declared no competing interest.

### Funding

- Swiss National Science Foundation: 10000473
- Swiss Cancer League: KFS-5422-08-2021-R
- Balgrist Foundation
- ResOrtho Foundation
- Swiss Government Excellence Scholarship: ESKAS-Nr: 2021.0468

