## Supplementary Information for "Chiron3D: an interpretable deep learning framework for understanding the DNA code of chromatin looping"

#### Supplementary Methods

**Raw data sources** We analyze CTCF HiChIP data from the Ewing sarcoma cell line A673_WT, published by Surdez et al [1]. Data were obtained from GEO (GEO accession number GSE133227). For the A673_WT cell line, there are a total of five reported replicates amounting to a total amount of read pairs of 320,219,396. For sequence coordinates, the reference genome hg19 was used. Cohesin and CTCF ChIP-seq profiles were also obtained from Surdez *et al.* (GEO: GSE133227). The A673 wildtype cohesin (SMC1A) HiChIP was obtained from Adane et al [2] (GEO: GSE165977).

**Reference genome** Windows of length 524,288 bp are tiled across the hg19 assembly with a stride of 50 kb. Each window later maps to a $105 \times 105$ contact matrix at 5 kb resolution. We exclude windows that cross chromosome ends or intersect with hg19 blacklist regions. This yields 59,812 candidate windows genome-wide before masking and 53,151 windows after our filtering.

**Preprocessing** The given CTCF HiChIP matrix was in .matrix.gz format and was converted to `.cool` using the `cooler` library in Python [3]. Here, the data for chromosome X and Y were excluded and the matrices log1p-transformed. Apart from that no additional matrix processing was applied. For the CTCF ChIP-seq data, we converted the given FASTQ files into BigWig files at a 1 base-pair resolution. The processing followed the steps outlined by Surdez et al. [1].

**Chiron3D Architecture Details** We initialized the sequence backbone using a pretrained Borzoi model [4, 5]. Having this foundation model as a sequence feature extractor allowed the sequence embeddings to be rich with genomic and epigenomic features relevant to regulatory elements and long-range chromatin interactions. Given a one-hot encoded DNA-sequence as input of shape $524{,}288 \times 4$, the backbone encodes the sequence via stacked 1D convolutional blocks (batch normalization, GELU, convolution) with strided pooling, reducing the resolution to 128 bp and producing a $4{,}096 \times 1{,}536$ feature map. An eight-layer multi-head self-attention module (8 heads) is then applied. A U-Net–style decoder [6] with skip connections restores the resolution to 32 bp, yielding embeddings of shape $16{,}384 \times 1{,}536$. Unlike in Borzoi, which center-crops to 196,608 bp for downstream track prediction, we retained the full 524,288 bp context and carry forward the complete $16{,}384 \times 1{,}536$ embedding to preserve long-range positional information relevant for modeling loop extrusion and boundary interactions.

We consider two adaptation regimes: (i) low-rank adaptation (LoRA) [7] applied to convolutional and linear layers as well as to the attention query and value projections (rank $r$=8); and (ii) a fully frozen backbone (no LoRA).

To map the 32 bp embeddings to the $105 \times 105$ contact target (525 kb at 5 kb resolution), we used three submodules. First, average pooling followed by a linear projection reduces the sequence to $105 \times 128$. Second, a transformer block stack (4 layers, 4 heads) models long-range dependencies at 5 kb resolution [8]. Third, we formed pairwise features by concatenating position-wise embeddings for all pairs, producing a $105 \times 105 \times 256$ tensor. A decoder with five dilated residual blocks (increasing dilation) aggregates multi-scale context, and a final $1 \times 1$ convolution emits the $105 \times 105$ prediction. The pairwise feature formation and dilated-residual decoder followed the C.Origami paradigm [9], reparameterized for a $105 \times 105$ target at 5 kb resolution, with upsampling replaced by pooling. Overall, the model has 185 M frozen parameters and either 2.5 M trainable parameters (adapter head only) or 4.6 M with LoRA.

**Evaluation Metrics** We evaluated the predicted log-transformed contact maps using Mean Squared Error (MSE), insulation score correlations, distance-stratified correlations, and the Structural Similarity Index Measure (SSIM). MSE was computed as the pixel-wise squared difference between the predicted and experimental matrices. To evaluate boundary predictions, we derived 1D insulation profiles for each map and computed their Pearson and Spearman (using scipy) correlations. Since contact frequencies naturally decay with genomic distance, we also calculated these correlations independently for each diagonal offset. Finally, to assess 2D structural fidelity, we used SSIM (using scikit-image). By measuring local structural covariance, SSIM evaluates the sharpness of topological features, where a higher SSIM value reflects better retention of the underlying structural features in the interaction maps. The SSIM data range was dynamically set to the minimum and maximum values of each matrix.

**Loop Datasets Construction** For the stripe symmetry analysis, we used the data provided by Surdez *et al.* that was obtained with the Tweed Algorithm [1]. For classification, we follow the rules outlined by [1] in their work. We calculated the ratio of the sum of the values for both stripes $s_x, s_y$, disregarding the 15 bins (75 kb) closest to the diagonal. Loops on the extremes of the asymmetry ratio were subsequently classified as asymmetrical. Through this, we ended up with 8,017 loop sites for CTCF HiChIP, 696 of which are asymmetrical, per our definition. Similarly, we obtained 2,251 loop sites for cohesin HiChIP, 101 of which are asymmetrical.

In order to obtain loops for the loop stability analysis, we used the FitHiChIP tool [10], which detected 26,130 CTCF HiChIP loops. However, for most of these loops, many neighboring loops were observed. Therefore, within regions of 50,000 bp, we retained only the loop with the lowest q-value, reducing the total number of CTCF loops to 12,273. In the next step, we removed all loops that intersected with the identified loops by the Tweed Algorithm [1], leaving us with 6,271 CTCF loops. For these loops, we checked whether at each loop boundary at least one cohesin subunit (RAD21, SMC1, STAG1, or STAG2) was present and whether the loop boundaries contained convergent CTCF motifs. This was done by using the CTCF forward and reverse PSSMs on the experimental CTCF peak regions at the loop boundaries, extracting the maximum binding affinity score. If the highest score at the left boundary corresponded to the forward CTCF motif and the highest score at the right boundary corresponded to the reverse CTCF motif, the loop was retained. The final number of CTCF loops was 1,895. Upon applying similar filtering for cohesin HiChIP loops, only 166 loops remain.

**Train Split** The final training/validation/test chromosome splits are: **train**: all autosomes except {5, 12, 13, 21, 2, 6, 19}, **validation**: {5, 12, 13, 21}, **test**: {2, 6, 19}.

**Model Training** Each data point consisted of a one-hot encoded DNA window of length $L_{\text{bp}} =$ 524,288 bp from the hg19 reference genome, together with its corresponding CTCF HiChIP contact submatrix at 5 kb resolution. Windows were tiled genome-wide with a 50 kb stride, mapping to $L = 105$ bins and a target matrix $M_{raw} \in \mathbb{R}_{\geq 0}^{L \times L}$. Targets were transformed with $M = \log\big(1 + M_{\text{raw}}\big)$. Training minimized the mean squared error on the log-transformed targets, $\mathcal{L}_{\text{MSE}} = \frac{1}{|\mathcal{I}|} \sum_{(i,j)\in\mathcal{I}} \big(\widehat{M}_{ij} - M_{ij}\big)^2$, where $\mathcal{I}$ indexes all pairs in the $L \times L$ window, $\widehat{M}$ is the prediction, and $M$ is the ground truth. Optimization was done using AdamW [11] (learning rate $5 \times 10^{-4}$, batch size 4) with mixed precision (bf16). A ReduceLROnPlateau [12] scheduler (factor 0.1, patience 5, minimum learning rate $1 \times 10^{-7}$) and early stopping using Pearson's correlation on the validation sets were used. Training converged in 24 hours on four RTX4090 GPUs. No test-time tuning was performed.

**Adaptation of the C.Origami model** We used C.Origami [9] as a strong baseline because it is state-of-the-art among contact map prediction models and easily adapts to our resolution. The original architecture predicted 10 kb Hi-C at 8,192 bp bins, upsampling a $210 \times 210$ matrix to $256 \times 256$. Our targets were 5 kb HiChIP measurements at $105 \times 105$, so we implemented the following as adaption: (i) We reduced the sequence and feature encoders by one layer. (ii) We halved the number of transformer blocks (8→4). (iii) We replaced the output upscaling with pooling to the desired $105 \times 105$ shape. We trained two C.Origami variants: DNA-only input and DNA+CTCF ChIP-seq (log-transformed) input. Because C.Origami operates on a 1 Mb field-of-view at 5 kb, its native output is $210 \times 210$. For head-to-head comparisons with Chiron3D, we center-cropped a $105 \times 105$ window and only evaluated against that.

We did not compare against the original C.Origami model, which required ATAC-seq as an additional input to avoid broader chromatin accessibility effects. Further, we also did not compare against DeepC or Orca, since these methods either require significant compute [13, 14] whereas Akita predicts observed-over-expected ($O/E$) contact maps, which are designed to emphasize structural features by regressing out the distance-decay relationship [15]. In contrast, Chiron3D targets CTCF HiChIP data, where the protein-mediated enrichment requires a model capable of capturing absolute interaction intensities and specific peak-to-background ratios that $O/E$ normalization may obscure.

**Attribution Scoring** Nucleotide-level importance was computed with input×gradient [16] in relation to the defined objectives *Asym* and *Stab*. These were then used as input for TF-MoDISco [17], which identifies recurrent predictive sequence patterns directly from attribution maps rather than from sequence enrichment alone. Briefly, TF-MoDISco extracts short high-scoring subsequences ("seqlets") from regions with strong contribution, partitions them into metaclusters according to the sign of their contribution, aligns similar seqlets, and aggregates them into consolidated motif representations. Here, positive and negative metaclusters denote seqlets whose contributions increase or decrease the selected objective, respectively. For visualization, related TF-MoDISco motifs were further grouped into broader motif clusters following the grouping from [18], and the reported seqlet share for a given loop type and boundary denotes the fraction of seqlets in that subset assigned to the corresponding motif cluster. Beyond visualization, the ratios *Asym* and *Stab* serve as scalar objectives for edit proposal: within a given loop window, we optimize candidate nucleotide edits to monotonically increase or decrease *Asym* or *Stab* as specified by the task. Concretely, to favor extrusion-like, stripe-dominated states we minimize *Stab*, whereas to promote stable loops we maximize *Stab*; stripe enhancement and reduction are achieved by maximizing and minimizing *Asym*, respectively.

**Edit Design** In silico edits were generated with Ledidi [19], , which formulates genomic edit design as an optimization problem in which the predictor is held fixed and the input sequence is modified to achieve a desired model output. Starting from the original one-hot encoded DNA sequence $X_0 \in \{0,1\}^{L_{\text{bp}}\times 4}$ , Ledidi relaxes the discrete sequence to a differentiable representation $g(W) = \text{softmax}(W/\tau)$ where $W$ is initialized from $X_0$ and $\tau > 0$ is the temperature. Given predictor $f(\cdot)$ and target objective value $\hat{y}$, we solve

$$\min_W \|g(W) - X_0\|_1 \;+\; \lambda \|f(g(W)) - \hat{y}\|_2^2,$$

where the $\ell_1$ term encourages few edits and the second term drives the desired change in model output thus balancing sparsity and task specific objective. In our application, $f(\cdot)$ is the frozen Chiron3D-based downstream objective wrapper and $\hat{y}$ specifies the requested increase or decrease in *Asym* or *Stab*, depending on the task. After convergence, a greedy pruning pass removes superfluous substitutions to yield a minimal discrete edit set. Ledidi also supports input masks to disallow edits in specified regions.

**Edit Labeling** Edits were considered unlabeled unless they overlapped a CTCF ChIP-seq peak or exhibited a change in CTCF motif affinity. CTCF binding affinity was quantified using the PSSM derived from the JASPAR MA0139.1 CTCF frequency matrix [20, 21]. Let the CTCF motif length be $L_C$=19, and define a local sequence window of $L_W$=41 bp centered on each edit (20 bp upstream and 20 bp downstream). For a window sequence $x_{1:L_W}$ (uppercase $\{A, C, G, T\}$), we construct a log-odds PSSM $M \in \mathbb{R}^{4\times L_C}$ assuming a uniform background $p_b$=0.25, with rows ordered as $\{A, C, G, T\}$. The log-odds score of a sequence segment $s$ of length $L_C$ is defined as

$$\text{score}(s; M) \;=\; \sum_{i=1}^{L_C} M_{\text{BASE2IDX}(s_i),\, i}$$

For each edit location, the maximum score is taken as the window-level CTCF affinity. The total change in PSSM is given by $\Delta\text{PSSM}_{\text{CTCF}} = \text{score}_{\text{post}} - \text{score}_{\text{pre}}$ where, $\text{score}_{\text{post}}$ is the maximum CTCF PSSM score computed on the edited DNA sequence and $\text{score}_{\text{pre}}$ is the corresponding maximum score on the original sequence.

[2] Biniam Adane, Gabriela Alexe, Bo Kyung A Seong, Diana Lu, Elizabeth E Hwang, Denes Hnisz, Caleb A Lareau, Linda Ross, Shan Lin, Filemon S Dela Cruz, et al. Stag2 loss rewires oncogenic

and developmental programs to promote metastasis in ewing sarcoma. *Cancer cell*, 39(6):827–844, 2021.

[8] Ashish Vaswani, Noam Shazeer, Niki Parmar, Jakob Uszkoreit, Llion Jones, Aidan N Gomez, Łukasz Kaiser, and Illia Polosukhin. Attention is all you need. *Advances in neural information processing systems*, 30, 2017.

[9] Jimin Tan, Nina Shenker-Tauris, Javier Rodriguez-Hernaez, Eric Wang, Theodore Sakellaropoulos, Francesco Boccalatte, Palaniraja Thandapani, Jane Skok, Iannis Aifantis, David Fenyö, et al. Cell-type-specific prediction of 3d chromatin organization enables high-throughput in silico genetic screening. *Nature biotechnology*, 41(8):1140–1150, 2023.

[10] Sourya Bhattacharyya, Vivek Chandra, Pandurangan Vijayanand, and Ferhat Ay. Identification of significant chromatin contacts from hichip data by fithichip. *Nature communications*, 10(1):4221, 2019.

[11] Ilya Loshchilov and Frank Hutter. Decoupled weight decay regularization. In *International Conference on Learning Representations*.

[17] Avanti Shrikumar, Katherine Tian, Žiga Avsec, Anna Shcherbina, Abhimanyu Banerjee, Mahfuza Sharmin, Surag Nair, and Anshul Kundaje. Technical note on transcription factor motif discovery from importance scores (tf-modisco) version 0.5. 6.5. *arXiv preprint arXiv:1811.00416*, 2018.

[18] Jeff Vierstra, John Lazar, Richard Sandstrom, et al. Global reference mapping of human transcription factor footprints. *Nature*, 583:729–736, 2020.

[19] Jacob Schreiber, Franziska Katharina Lorbeer, Monika Heinzl, Yang Young Lu, Alexander Stark, and William Stafford Noble. Programmatic design and editing of cis-regulatory elements. *bioRxiv*, pages 2025–04, 2025.

[20] Ieva Rauluseviciute, Rafael Riudavets-Puig, Romain Blanc-Mathieu, Jaime A Castro-Mondragon, Katalin Ferenc, Vipin Kumar, Roza Berhanu Lemma, Jérémy Lucas, Jeanne Chèneby, Damir Baranasic, et al. Jaspar 2024: 20th anniversary of the open-access database of transcription factor binding profiles. *Nucleic acids research*, 52(D1):D174–D182, 2024.

[21] Gary D Stormo. Dna binding sites: representation and discovery. *Bioinformatics*, 16(1):16–23, 2000.

### Supplementary Figures

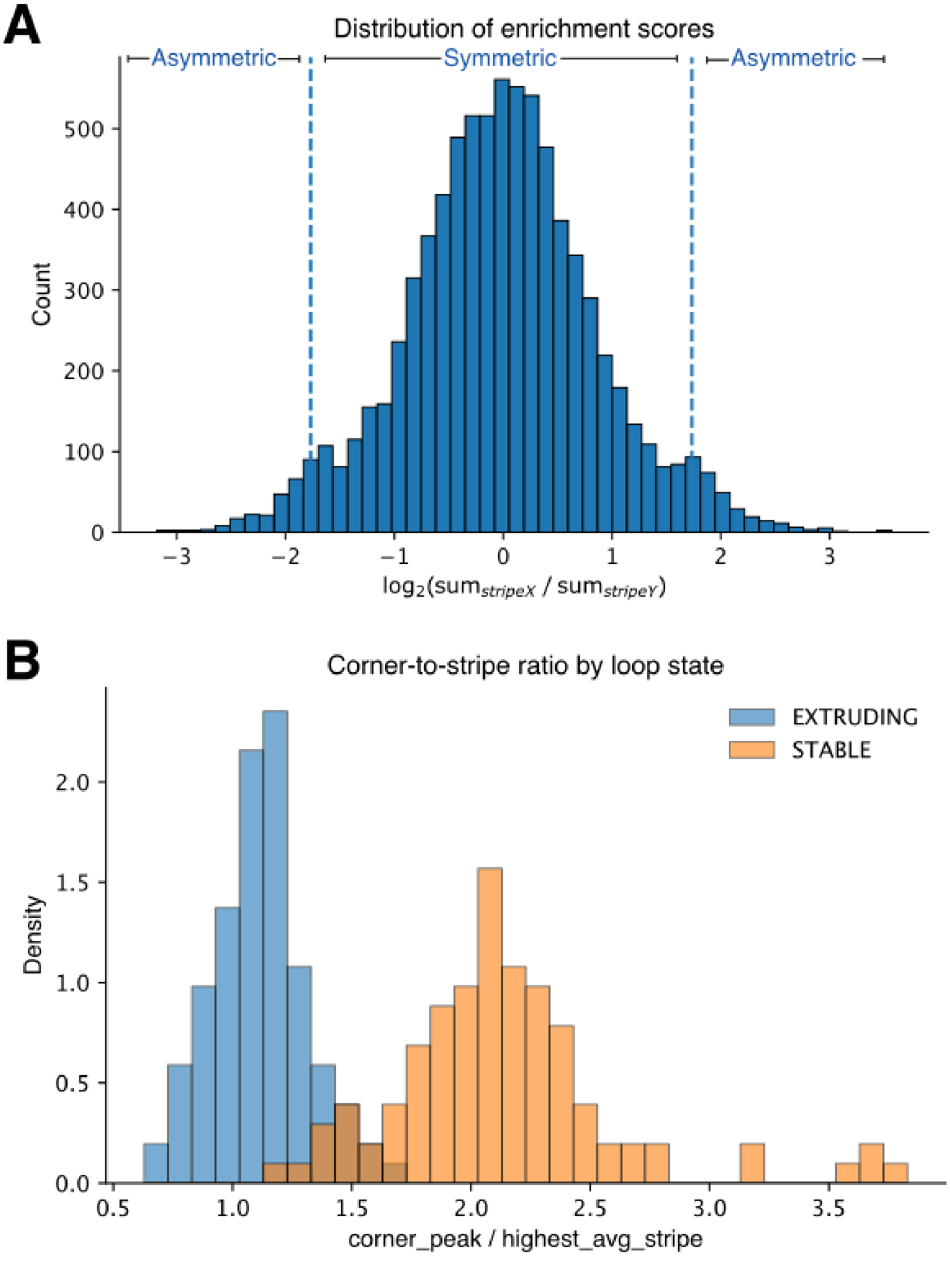


Figure S1: **Distributions of different loop types across our defined scoring metrics. A,** Distribution of Tweed-called loops across stripe ratio between X and Y stripes. The top and bottom 5% of the loops are labeled as asymmetric and otherwise symmetric. **B,** The distribution of ratio of corner interaction and highest average stripe intensity (defined by us as *Stab* score) across loops labelled as Stable and Extruding.

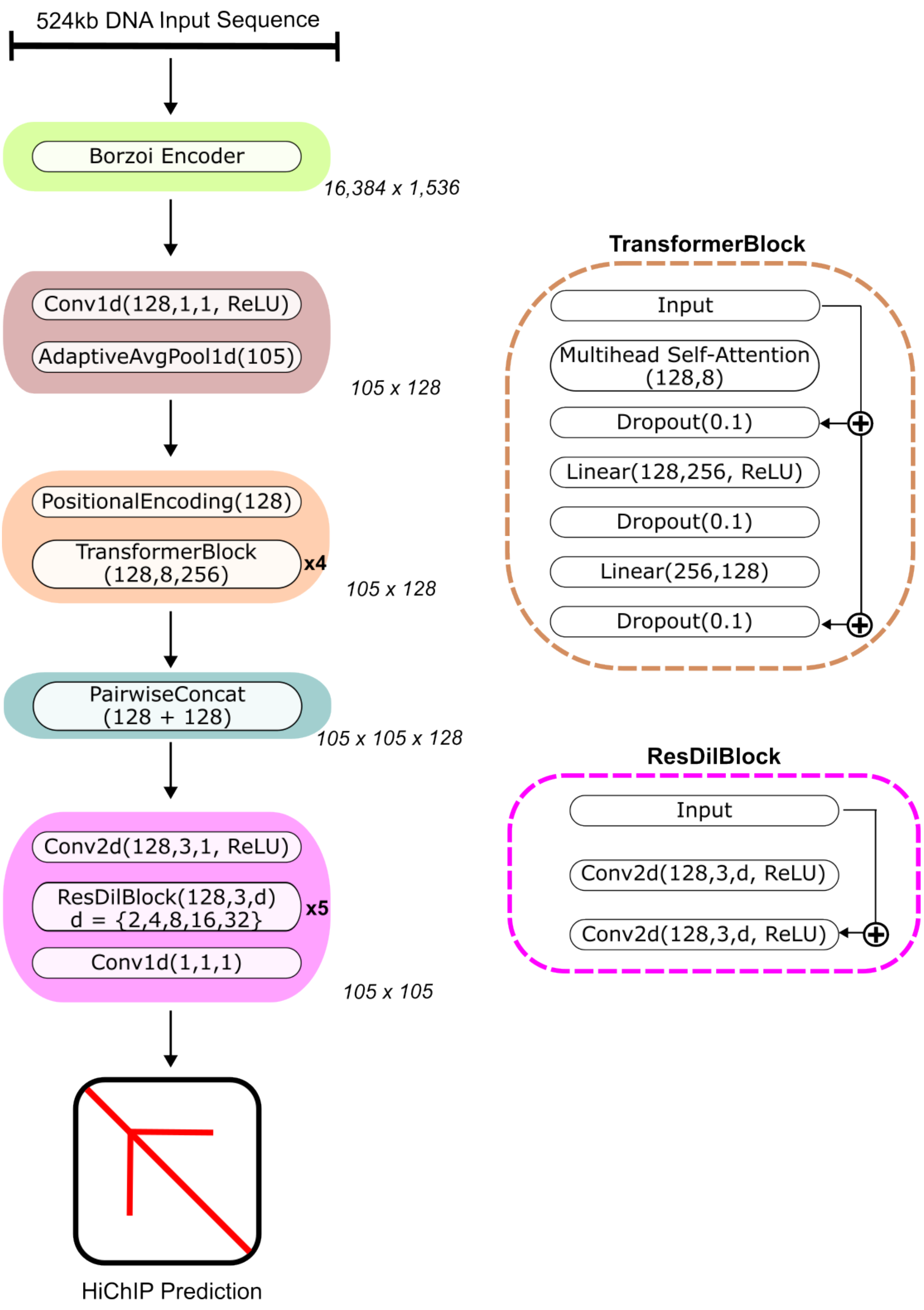


Figure S2: **Detailed architecture diagram of Chiron3D.** Chiron3D consists of a pretrained Borzoi sequence backbone that encodes the input DNA sequence into contextual embeddings capturing regulatory and long-range genomic features. These embeddings are aggregated to the target resolution and processed with transformer layers to model long-range dependencies. Pairwise combinations of positional embeddings represent potential chromatin contacts, and a dilated residual decoder predicts the final contact map. The backbone is used either frozen or adapted using low-rank adaptation (LoRA).

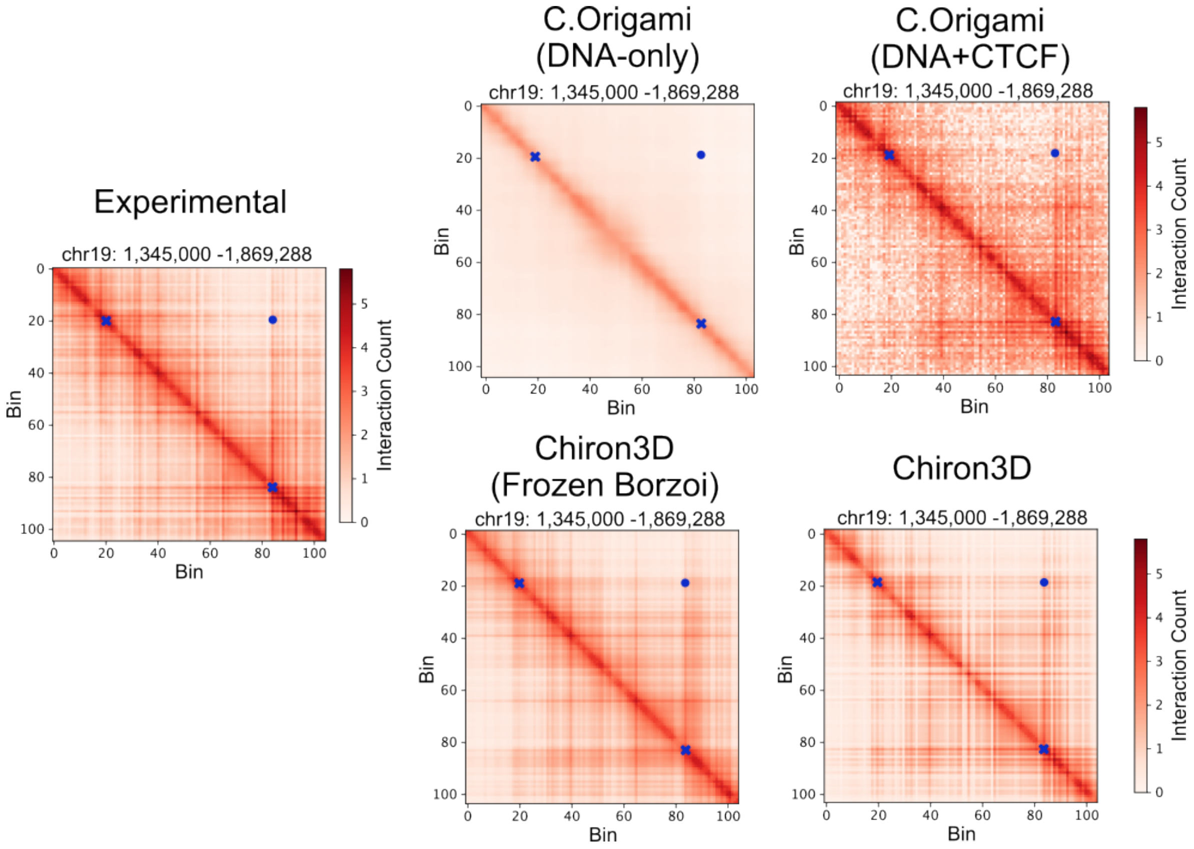


Figure S3: **Qualitative comparison between model predictions and experimental HiChIP at a representative test locus.** All models are compared on test locus chr19:1,345,000-1.869,288. The loop boundaries are marked with blue crosses whereas the corner interaction is depicted with a blue dot.

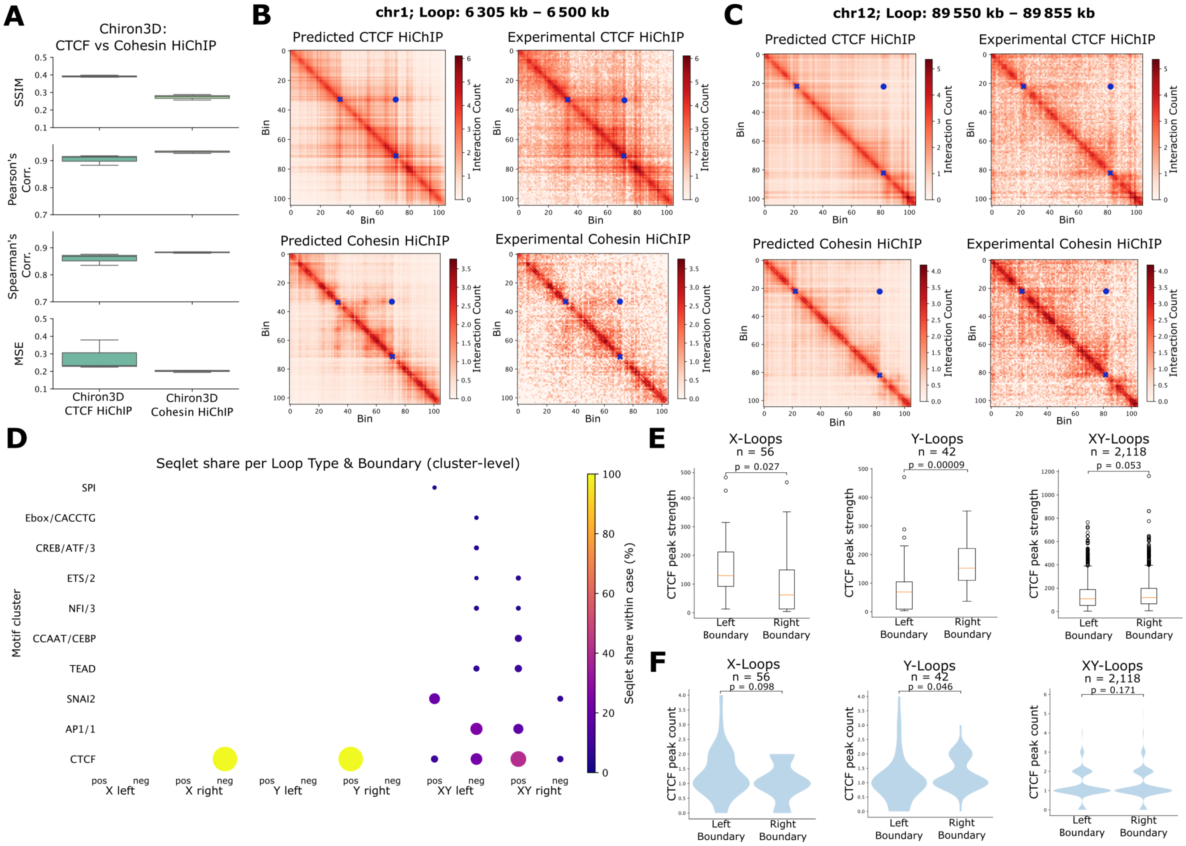


Figure S4: **CTCF is a driving factor for defining loop symmetry in cohesin HiChIP. A,** Performance of Chiron3D is compared between two training datasets: CTCF HiChIP and cohesin (SMC1A) HiChIP data. Model performance is calculated using Structural similarity index measure (SSIM), Pearson's correlation, Spearman's correlation, and mean squared error (MSE). **B-C,** Outputs of the two Chiron3D models are compared against experimental HiChIP contact maps for two representative loci. **D,** Major transcription factor motifs identified using TF-MoDISco on boundary regions of extruding cohesin loops. **E,** Measurements of CTCF ChIP-seq signal at loop boundaries of X- and Y-asymmetric cohesin loops and symmetric cohesin loops. $n$ is the number of loops of each type. **C**, Number of CTCF ChIP-seq peaks at loop boundaries of X- and y-asymmetric cohesin loops and symmetric cohesin loops. All p-values are calculated using Wilcoxon signed-rank test.

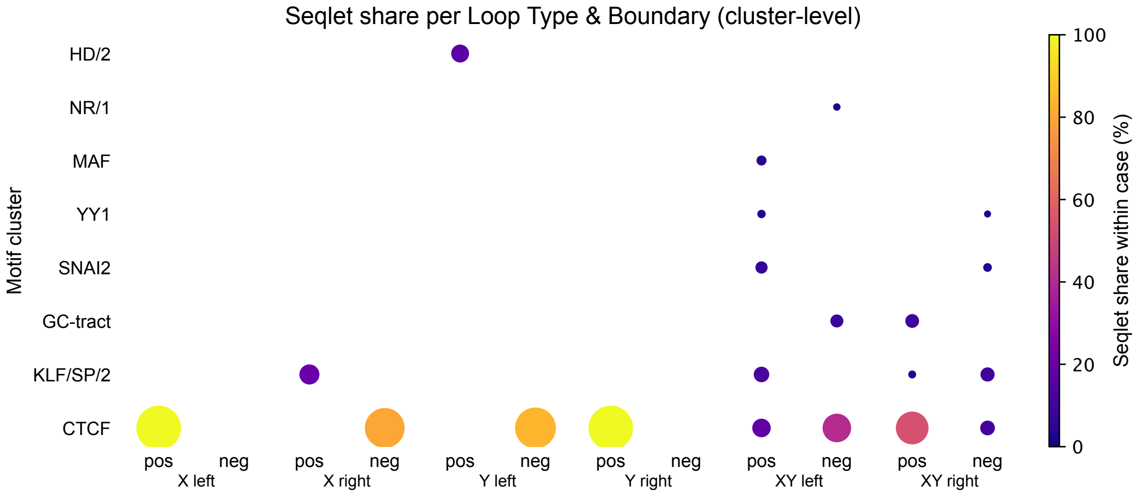


Figure S5: **CTCF as most important motif detected for loop anchorage strength.** Filtered and aggregated results from TF-MoDISco analysis on boundary regions of extruding loops.

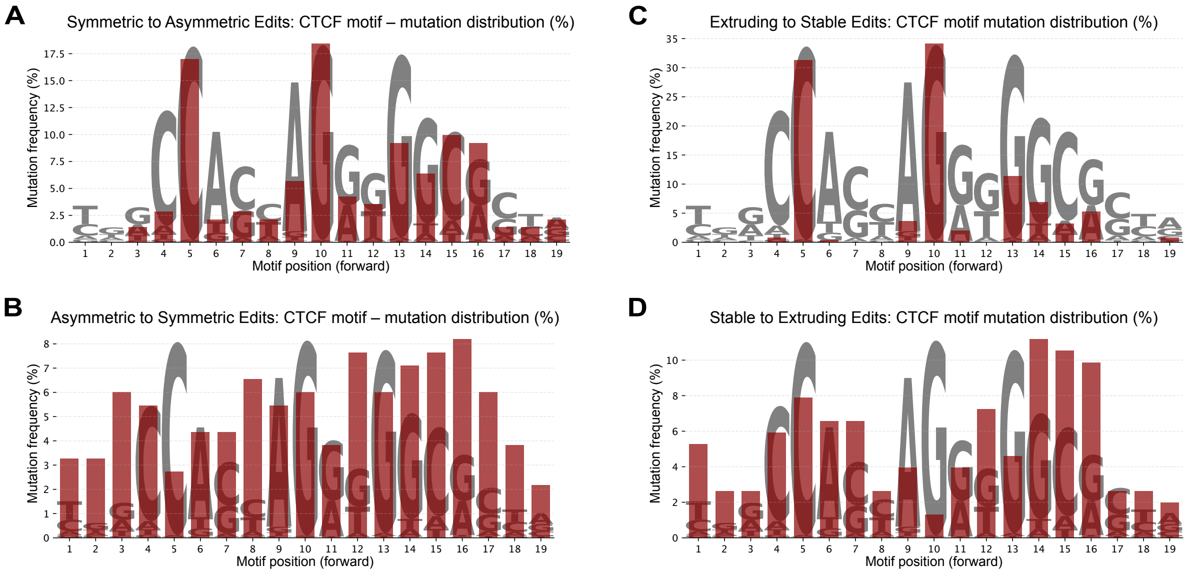


Figure S6: **Distribution of edits that fall within CTCF motif for all different objectives.** **A**, Distribution of edits within CTCF motif when proposing edits to make a loop more asymmetric. **B**, Distribution of edits within CTCF motif when proposing edits to make a loop more symmetric. **C**, Distribution of edits within CTCF motif when proposing edits to make a loop more stable. **D**, Distribution of edits within CTCF motif when proposing edits to make a loop more extruding.

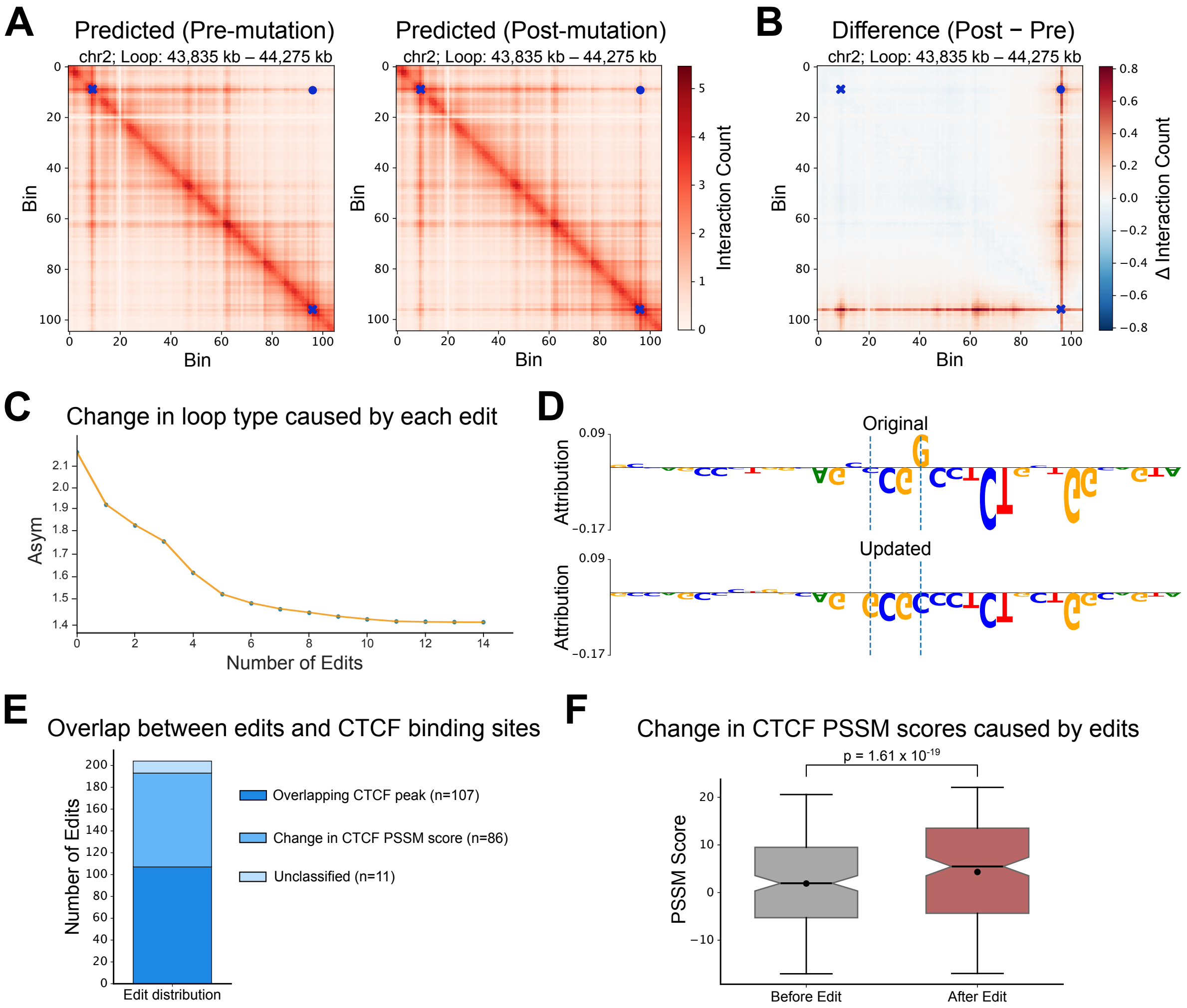


Figure S7: **Higher CTCF anchoring strength and more CTCF motifs increase anchorage probability.** **A**, Example of a region from the test set where two edits within single motif are proposed to make a asymmetric loop more symmetric. **B**, The difference in predicted contact maps caused by the proposed edit. **C**, Line plot showing change in the optimization function ($Asym$) as a function of number of proposed edits. **D**, Attribution scores before and after the edit was introduced in the sequence. **E**, Proportion of edits, proposed for symmetric loops in the test dataset, that overlap with CTCF regions. **F**, Change in CTCF PSSM scores caused by the edits across all symmetric loops in the test data. Significance is calculated using Wilcoxon signed-rank test.

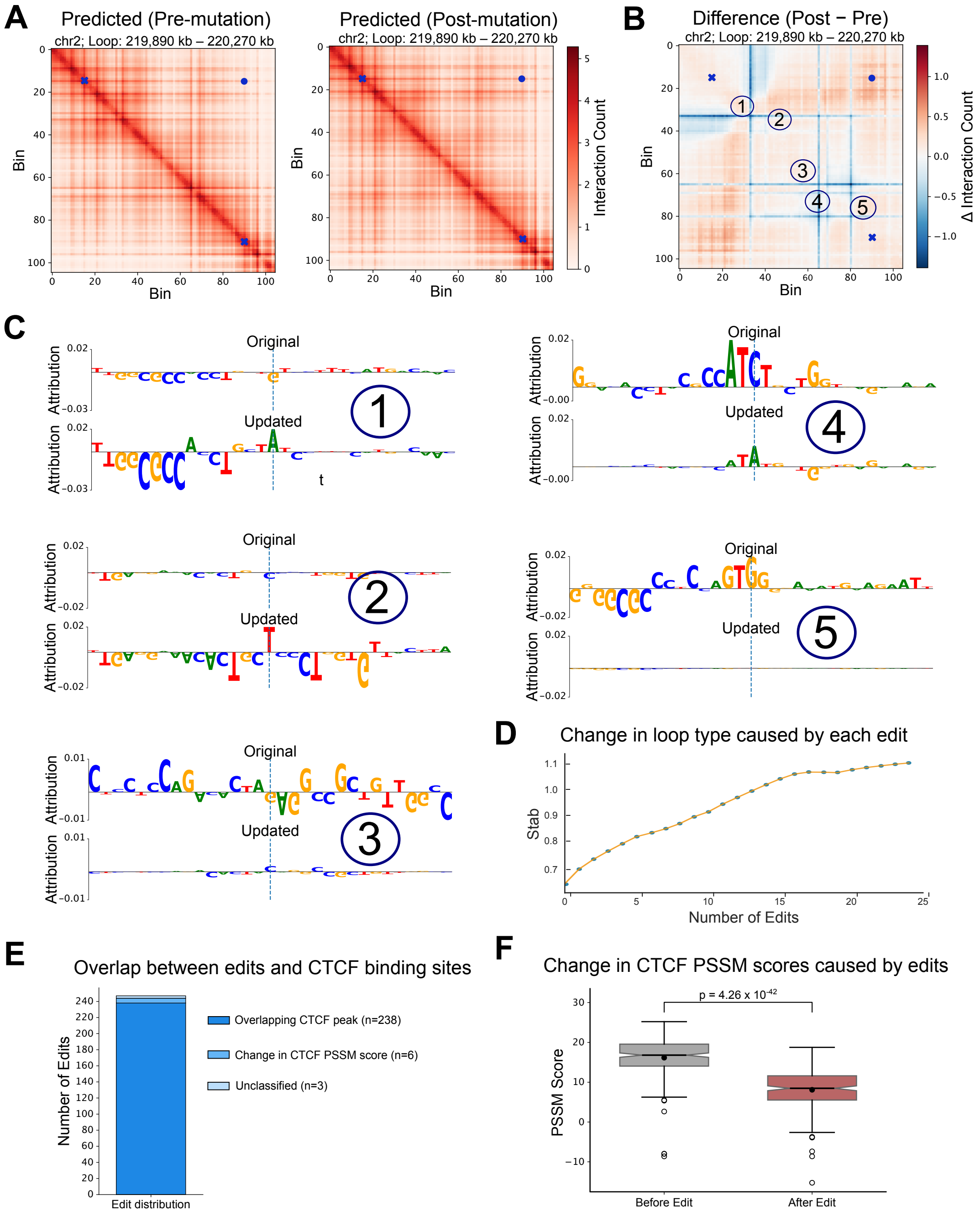


Figure S8: **Fewer intra-loop CTCF sites lead to more stable loop domain.** **A**, Example of a region from the test set where total of five edits are proposed to make a loop more stable. **B**, The difference in predicted contact maps caused by all five proposed edits. **C**, Attribution scores before and after each edit was introduced in the sequence. **D**, Line plot showing change in the optimization function (*Stab*) as a function of number of proposed edits. **E**, Proportion of edits, proposed for symmetric loops in the test dataset, that overlap with CTCF regions. **F**, Change in CTCF PSSM scores caused by the edits across all symmetric loops in the test data. Significance is calculated using Wilcoxon signed-rank test.

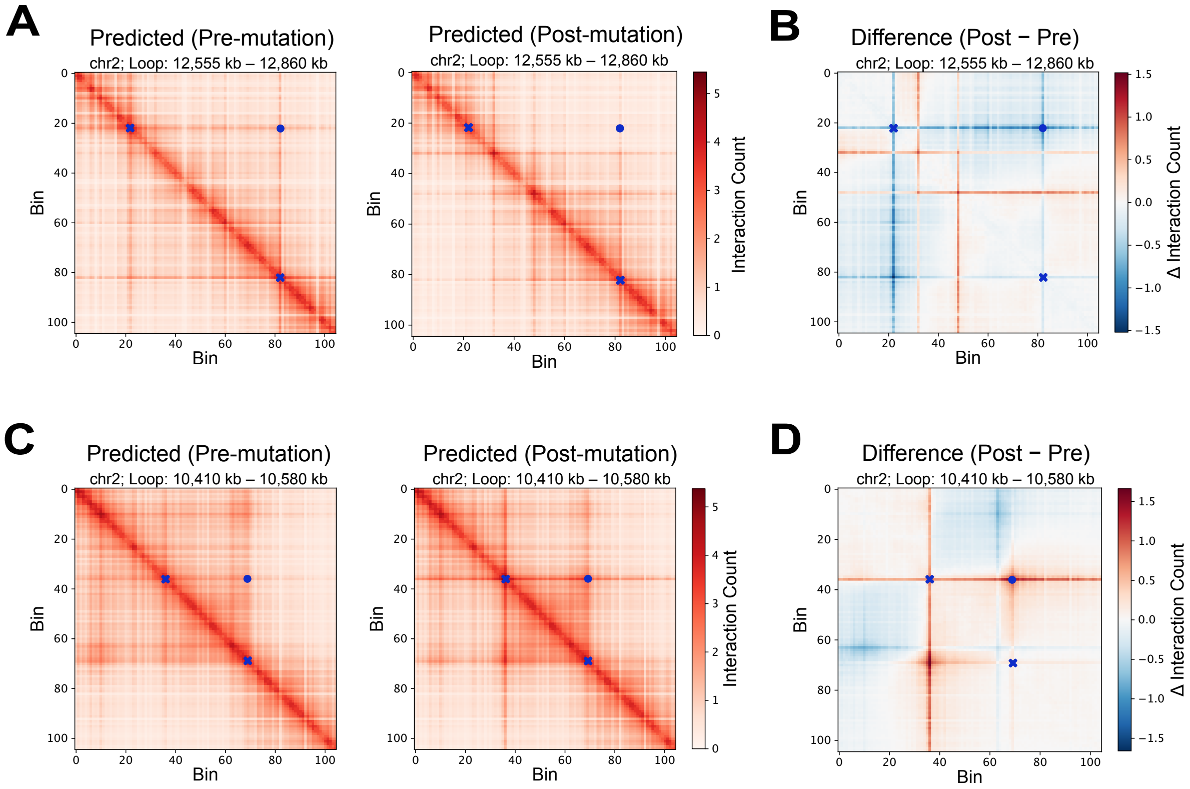


Figure S9: **Detected antipattern on boundary regions if no masking is applied.** **A**, Example of a region from the test set where additional edits on the boundaries are proposed to make a loop more stable. **B**, The difference in predicted contact maps caused by the proposed edits. **C**, Example of a region from the test set where additional edits on the boundaries are proposed to make a loop more extruding. **D**, The difference in predicted contact maps caused by the proposed edits.

### Supplementary Tables

Table S1: Edits predicted by our framework to alter loop anchoring and stripe symmetry overlap with known SNPs.

| Chrom | Position | SNP ID | Change in Loop Type |
|---|---|---|---|
| chr19 | 14184412 | rs1352588834 | Asymmetric to Symmetric |
| chr19 | 15272625 | rs961480933 | Asymmetric to Symmetric |
| chr19 | 15273396 | rs1403803265 | Asymmetric to Symmetric |
| chr19 | 41933686 | rs566412227 | Asymmetric to Symmetric |
| chr19 | 56840222 | rs1339965976 | Asymmetric to Symmetric |
| chr2 | 103068666 | rs1233062627 | Asymmetric to Symmetric |
| chr2 | 174313763 | rs1406762974 | Asymmetric to Symmetric |
| chr2 | 236679937 | rs1463496071 | Asymmetric to Symmetric |
| chr6 | 11578350 | rs1360966452 | Asymmetric to Symmetric |
| chr6 | 20092481 | rs148095721 | Asymmetric to Symmetric |
| chr6 | 38941160 | rs571045563 | Asymmetric to Symmetric |
| chr6 | 41754317 | rs1369265027 | Asymmetric to Symmetric |
| chr6 | 166263711 | rs540744523 | Asymmetric to Symmetric |
| chr19 | 4186582 | rs1469806546 | Symmetric to Asymmetric |
| chr19 | 11593584 | rs752861056 | Symmetric to Asymmetric |
| chr19 | 15981885 | rs1423089126 | Symmetric to Asymmetric |
| chr19 | 49979697 | rs1463648878 | Symmetric to Asymmetric |
| chr19 | 52111363 | rs987138279 | Symmetric to Asymmetric |
| chr2 | 86448408 | rs749579 | Symmetric to Asymmetric |
| chr2 | 178539127 | rs985932703 | Symmetric to Asymmetric |
| chr6 | 29802794 | rs994405403 | Symmetric to Asymmetric |
| chr6 | 30646925 | rs1363038402 | Symmetric to Asymmetric |
| chr6 | 40296317 | rs989998410 | Symmetric to Asymmetric |
| chr6 | 55845615 | rs994144485 | Symmetric to Asymmetric |
| chr6 | 55845622 | rs1398978912 | Symmetric to Asymmetric |
| chr6 | 57178837 | rs1230360127 | Symmetric to Asymmetric |

Table S2: Edits predicted by our framework to alter loop stability overlap with known SNPs.

| Chrom | Position | SNP ID | Change in Loop Type |
|---|---|---|---|
| chr19 | 28963962 | rs1024259702 | Stable to Extruding |
| chr19 | 29132131 | rs990357913 | Stable to Extruding |
| chr19 | 29350516 | rs959578078 | Stable to Extruding |
| chr19 | 29360632 | rs1200260765 | Stable to Extruding |
| chr19 | 41488726 | rs1225557199 | Stable to Extruding |
| chr19 | 41497265 | rs770247931 | Stable to Extruding |
| chr19 | 41571108 | rs891638207 | Stable to Extruding |
| chr2 | 46363023 | rs1239033001 | Stable to Extruding |
| chr2 | 57644490 | rs1388982439 | Stable to Extruding |
| chr2 | 68022148 | rs1335644990 | Stable to Extruding |
| chr2 | 84300968 | rs1299983807 | Stable to Extruding |
| chr2 | 84517811 | rs572069114 | Stable to Extruding |
| chr2 | 173091262 | rs984242323 | Stable to Extruding |
| chr6 | 24596304 | rs377520480 | Stable to Extruding |
| chr6 | 24596338 | rs1172584687 | Stable to Extruding |
| chr6 | 24596361 | rs138139227 | Stable to Extruding |
| chr6 | 24596538 | rs755928541 | Stable to Extruding |
| chr6 | 24667267 | rs1396785046 | Stable to Extruding |
| chr6 | 51097101 | rs1043196577 | Stable to Extruding |
| chr6 | 72752943 | rs1253717685 | Stable to Extruding |
| chr6 | 72761593 | rs974958979 | Stable to Extruding |
| chr6 | 100094035 | rs1487895477 | Stable to Extruding |
| chr6 | 100116418 | rs890808485 | Stable to Extruding |
| chr6 | 105142621 | rs1299177991 | Stable to Extruding |
| chr6 | 110500832 | rs578166434 | Stable to Extruding |
| chr6 | 121427160 | rs748024905 | Stable to Extruding |
| chr6 | 122001377 | rs1280058946 | Stable to Extruding |
| chr6 | 124028935 | rs973702849 | Stable to Extruding |
| chr6 | 130938936 | rs1486587060 | Stable to Extruding |
| chr6 | 153795995 | rs1003036770 | Stable to Extruding |
| chr6 | 161920069 | rs1364562969 | Stable to Extruding |
| chr6 | 161920813 | rs890713448 | Stable to Extruding |
| chr19 | 1000246 | rs1335548967 | Extruding to Stable |
| chr19 | 1001836 | rs577021345 | Extruding to Stable |
| chr19 | 1626597 | rs1356861490 | Extruding to Stable |
| chr19 | 3327143 | rs933235088 | Extruding to Stable |
| chr19 | 10380296 | rs1248762287 | Extruding to Stable |
| chr19 | 10594423 | rs772359649 | Extruding to Stable |
| chr19 | 14613354 | rs970832964 | Extruding to Stable |
| chr19 | 39369651 | rs200035722 | Extruding to Stable |
| chr19 | 40970424 | rs962747189 | Extruding to Stable |

*Continued on next page*

| Chrom | Position | SNP ID | Change in Loop Type |
|---|---|---|---|
| chr19 | 45831508 | rs535218778 | Extruding to Stable |
| chr19 | 47019264 | rs1384204628 | Extruding to Stable |
| chr19 | 47735125 | rs535868842 | Extruding to Stable |
| chr19 | 49249663 | rs28400017 | Extruding to Stable |
| chr19 | 49249858 | rs1316792237 | Extruding to Stable |
| chr19 | 49729472 | rs570591062 | Extruding to Stable |
| chr19 | 49769797 | rs1350743534 | Extruding to Stable |
| chr2 | 11925230 | rs1169425282 | Extruding to Stable |
| chr2 | 73089842 | rs1422759286 | Extruding to Stable |
| chr2 | 85659257 | rs995889710 | Extruding to Stable |
| chr2 | 220341168 | rs1427008570 | Extruding to Stable |
| chr2 | 220406445 | rs1199091122 | Extruding to Stable |
| chr2 | 241535262 | rs565982387 | Extruding to Stable |
| chr6 | 3404354 | rs1182997219 | Extruding to Stable |
| chr6 | 31335134 | rs1473229512 | Extruding to Stable |
| chr6 | 31430752 | rs3132090 | Extruding to Stable |
| chr6 | 37027188 | rs1427730174 | Extruding to Stable |
| chr6 | 39271138 | rs1329248215 | Extruding to Stable |
